# Impact of a tilted coverslip on two-photon and STED microscopy

**DOI:** 10.1101/2023.10.26.564142

**Authors:** Guillaume Le Bourdelles, Luc Mercier, Johannes Roos, Stephane Bancelin, U. Valentin Nägerl

## Abstract

The advent of super-resolution microscopy has opened up new avenues to unveil brain structures with unprecedented spatial resolution in the living state. Yet, its application to live animals remains a genuine challenge. Getting optical access to the brain *in vivo* requires the use of a ‘cranial window’, whose mounting greatly influences image quality. Indeed, the coverslip used for the cranial window should lie as orthogonal as possible to the optical axis of the objective, or else significant optical aberrations occur. In this work, we assess the effect of the tilt angle of the coverslip on STED and two-photon microscopy, in particular image brightness and spatial resolution. We then propose an approach to measure and reduce the tilt using a simple device added to the microscope, which can ensure orthogonality with a precision of 0.07°.

## 1. Introduction

The emergence of super-resolution microscopy has revolutionized the field of bio-imaging, providing researchers with the ability to visualize and study cellular structures and processes at a spatial scale that had been out of reach for light microscopy [1]. Over the years, various superresolution approaches have been demonstrated to overcome the diffraction limit of light and have found applications in numerous fields, such as cell biology, neuroscience, immunology [2,3], as well as polymer [4] and material [5] sciences.

Among these techniques, Stimulated Emission Depletion (STED) Microscopy [6] stands out as a laser-scanning technique, where the point-spread-function (PSF) of the microscope is physically improved. It utilizes a combination of two laser beams at different wavelengths: an excitation beam and a depletion beam. The excitation beam is focused onto the sample, causing fluorescence of the labeled molecules in the focal region. Concomitantly, the depletion beam is used to induce stimulated emission of fluorescence from the excited molecules. By spatially shaping the depletion beam as a *donut* or *bottle* pattern and overlapping it with the excitation beam, it is possible to deactivate the excited fluorophores in the outer regions of the focal volume, effectively confining the emission to a sub-diffraction spot, typically down to tens of nanometers [7].

Compared to other super-resolution approaches, STED microscopy enables fast imaging and offers relatively high penetration depth in scattering samples, making it suitable for live-tissue imaging [8–12]. As the technology keeps progressing, new variations and technical improvements push the boundaries of resolution [13] and depth penetration [14,15], opening new biological systems to nanoscale investigation [16,17]. Notably, several groups have pioneered the application of STED microscopy in living organisms to study neuronal connectivity, synaptic dynamics and protein localization in the mouse brain *in vivo* [18–23]. This typically involves a surgical procedure to implant a cranial window, where a small part of the skull is replaced with a glass coverslip to provide optical access to the brain [24]. Yet, performing STED microscopy *in vivo* is a much greater challenge than imaging fixed samples or isolated cells. Indeed, STED critically depends on the spatial shaping of the depletion beam, which can be strongly affected by any heterogeneity in the optical path (both in the microscope or in the sample). This ultimately determines the quality of the final image [10,14,25,26].

In optical microscopy, a coverslip is typically placed on the sample to protect it and provide a flat surface for imaging. As the last optical element before entry into the sample, the horizontality and flatness of this coverslip is of utmost importance to maintain the quality of the excitation and depletion PSFs. In an inverted microscope, which is the predominant design for STED microscopy, the sample is viewed from below through the coverslip, typically using an oil immersion objective. In this configuration, there is little refractive index mismatch between the immersion media and the coverslip. In combination with index-matching embedding media [27], this configuration generates minimal distortion of the PSF. Moreover, because of the inverted design, the orthogonality of the coverslip to the optical axis is largely ensured by the microscope stage, which facilitates the use of other immersion media (e.g., glycerin, silicone oil, water).

Yet, for *in vivo* imaging, oil objectives are not an option because of their short working distance, and neither is the use of inverted microscopes. Rather, water immersion objectives are generally used, leading to a significant refractive index mismatch with the coverslip. Additionally, chances are - unless appropriate technical measures are taken - that the optical axis is not orthogonal to the coverslip of the cranial window. Depending on this degree of non-orthogonality, the PSFs may get severely distorted, leading to signal loss and blurry images that will compromise STED performance.

In this work, we numerically and experimentally characterize the effects of a tilted coverslip on the different PSFs involved (excitation, depletion and effective fluorescence) for two different objectives (water and silicone oil immersion). We then propose and validate a straightforward approach to measure and correct this tilt directly under the microscope objective as a way to remedy the problem.

## 2. Material and Methods

### 3.1 PSF computation

The PSFs of the excitation and STED beams were calculated using a custom-written Matlab script based on the vectorial diffraction theory introduced by Richard and Wolf [28,29]. This approach allows to determine the electromagnetic field in the focal region of a high numerical aperture objective (NA ≥ 0.7). To take into account the influence of planar dielectric interfaces, we used the formalism developed by Törok *et al*. [30]. The effective fluorescence PSF (I_eff_) was deduced from the excitation (I_2P_) and STED PSF (I_STED_).

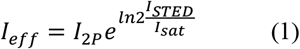

where I_sat_ is the saturation factor, which models the efficiency of the stimulated emission depletion [31]. In our numerical simulations, this parameter was empirically adjusted so that the full-width at half maximum (FWHM) of the effective PSF fits the one experimentally measured on our microscope. For a complete description of the calculation see [23].

The impact of a stratified medium on the focusing of light has been extensively studied [32–34]. In addition, the case of focusing with high NA through a tilted dielectric surface has been analytically derived [35,36]. Yet, surprisingly little effort has been undertaken to characterize numerically and experimentally the effect of a coverslip tilt angle on the PSF of the microscope. To the best of our knowledge, only a single research paper addresses this effect [37] using an undisclosed algorithm. In our code, instead of tilting the coverslip by an angle γ, which is analytically complex, we used an elegant solution proposed by S. Berning [38], which transforms the pupil function to model the tilt of the objective by an angle of −γ (Fig. 1.a). In this case, since no assumption is made regarding the symmetry of the pupil function for the calculation of the focal field, the integral for the propagation is the same as in the non-tilted case. As depicted in (Fig. 1.b), the new pupil function Aγ(θ,φ) corresponds to a rotation of the initial spherical wavefront A(θ,φ) around the origin O, followed by an orthographic projection back on the pupil plane. For the PSF calculation, the integration angle is then expanded from α (corresponding to the NA of the objective) to α+|γ|, while the part of the spherical wavefront falling out of the pupil function is set to 0 (green cone in Fig. 1.b).

**Fig. 1.**
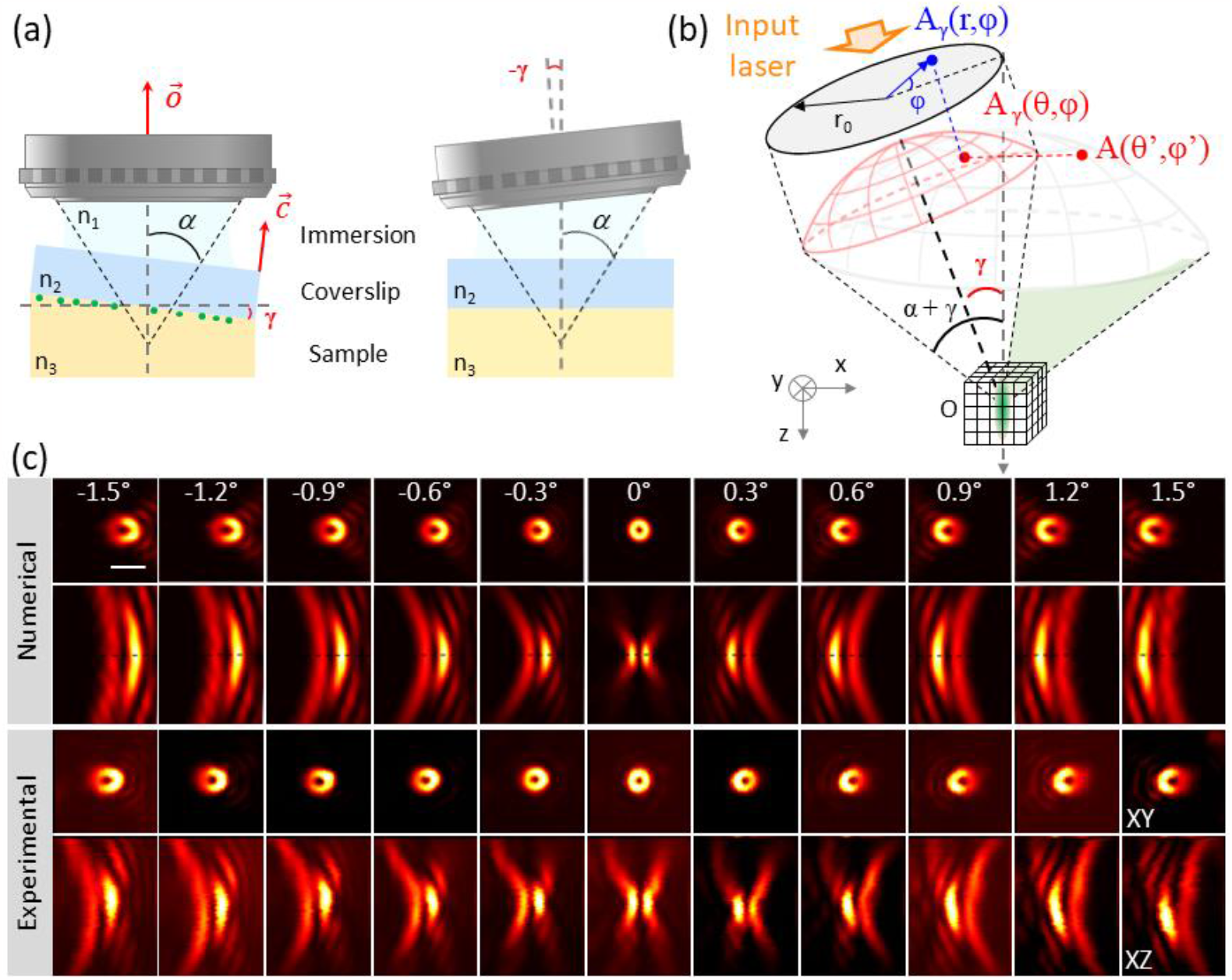
(a) Schematic representation of the simulations illustrating the different interfaces. Instead of tilting the coverslip, it is equivalent to rotate the objective by an angle −γ. (b) This can simply be modeled through a modification of the pupil function. The light waves are then focused by a high NA objective lens and propagated through a stratified medium composed of the immersion medium, the coverslip and the sample, to determine the PSF in the vicinity of the focus. (c) Numerical and experimental STED PSFs, highlighting the impact of a tilted coverslip on the *donut*-shape. Scale bar, 1 μm.

### 3.2 Experimental procedure

Imaging was performed using a custom-built upright 2P-STED microscope as previously described (see [10,19,23] for a complete description). In all the following, 2D-STED and z-STED refer to images acquired using pure *donut* or pure *bottle* beams, respectively. Gold bead samples were used to assess the *donut* or *bottle* beam quality, while optical resolution was assessed by imaging fluorescent beads immobilized on a glass coverslip. Samples were prepared with either gold beads (150 nm Gold nanospheres, Sigma-Aldrich) or fluorescent beads (yellow-green fluorescent beads, 170 nm in diameter, Invitrogen) placed on polylysine-coated #1 coverslips. Samples were left to dry overnight and then transferred to a microscope glass slide, where they were sealed with nail polish. In order to evaluate the effect of the coverslip tilt angle, samples were placed on a tiltable mechanical stage (TTR001/M, Thorlabs), enabling us to vary the tilt angle relative to the microscope objective axis in a precise and systematic way. STED PSFs were originally checked and optimized at a tilt angle of 0 ± 0.03°.

## 3. Results

### 3.1 Impact of coverslip tilt on STED microscopy performance

We first evaluated the effect of the tilt on the STED PSFs, both for donut and bottle beams. To that end, we placed the stage under the objective and introduced a controlled tilt angle between the optical axis of the objective 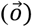 and the normal vector of the coverslip 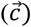. We performed measurements over a range of −1.5° to1.5° with 0.3° steps. Experimental results are displayed in Fig. 1.c together with the corresponding numerically simulated PSFs, clearly illustrating the impact of the tilt angle on the symmetry of the PSF, as well as on the null in the center. As expected, the aberrations introduced by a tilted coverslip are dominated by coma. To check the reproducibility of our measurements, we also evaluated the PSFs obtained when tilting the coverslip from 0° to 1.5° then to −1.5° and back to 0° with 0.3° steps (see Supp Fig.1 and 2 for donut and bottle beams, respectively).

Although informative, knowing the STED PSF is not sufficient to predict the quality of the final fluorescence image. To do so, we measured and computed the fluorescence PSFs as a function of tilt angle using fluorescent beads. Figure 2 displays the experimental effective PSFs in lateral (XY) and axial (XZ) directions without STED beam (2P) and in 2D-STED and z-STED modes, measured on different fluorescent beads. Additionally, we performed the corresponding numerical simulations (see Supp. Fig. 3). As expected, the distortions of the STED PSF observable in Fig. 1.c lead to strongly degraded effective fluorescence PSFs. As the fluorescent beads were relatively large (170 nm in diameter), their images are inevitably wider than the actual lateral resolution of our STED microscope (about 70 nm [10]). While smaller beads approximate more closely a fluorescent point source, they suffer from lower signal and increased photobleaching, making them impractical for the evaluation of the resolution at higher tilt angles (≥ 1°).

**Fig. 2.**
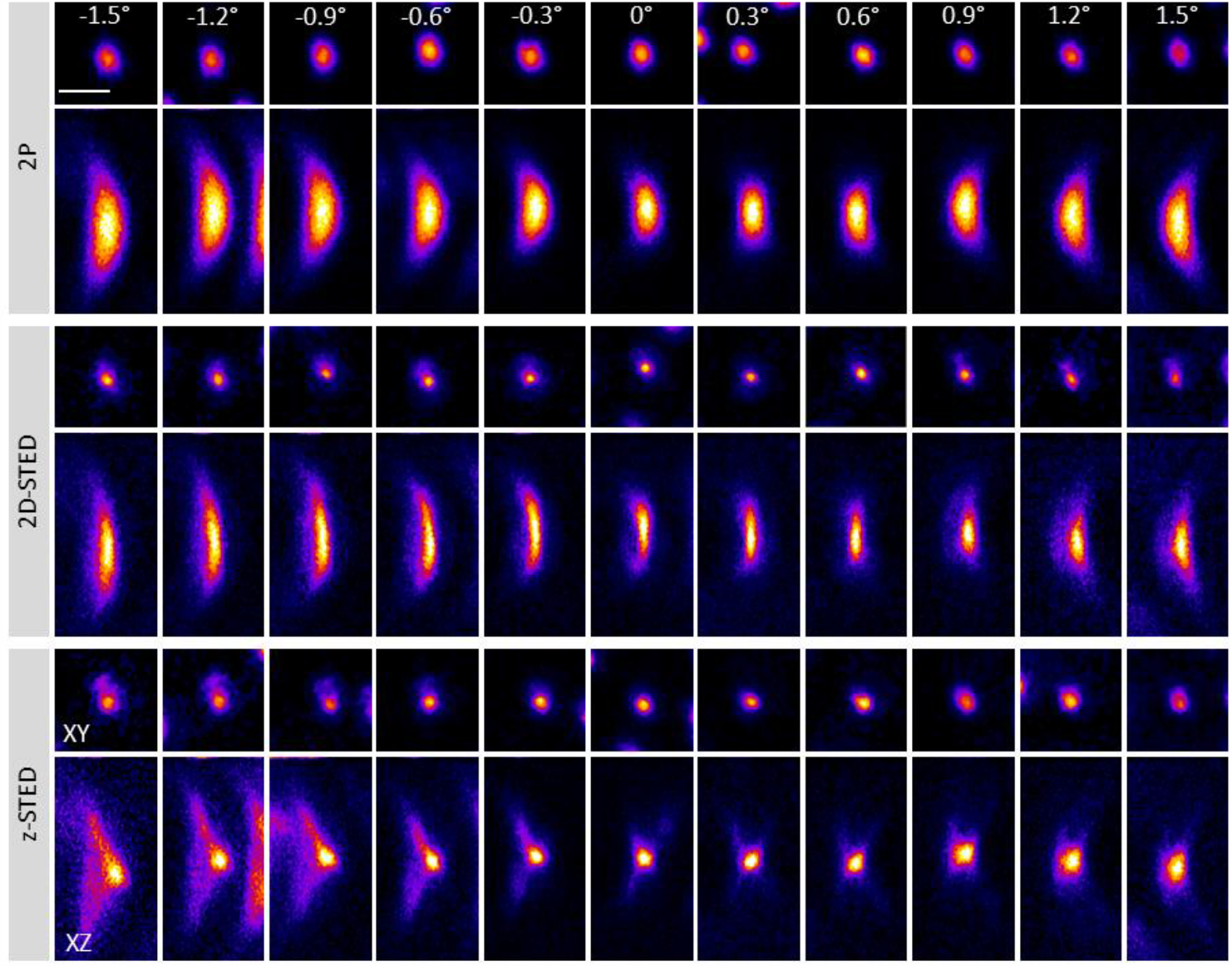
Impact of the coverslip tilt on the effective PSF measured on fluorescent beads in 2P, 2D-STED and z-STED modes. Scale bar, 1 μm. Look-up table is adjusted for every image to better highlight change of size and symmetry. Normalized intensities are plotted in Fig. 3.

The distortion of the PSF arises from the different refractive indices at the two interfaces ‘immersion medium-glass’ and ‘glass-sample’ (see Fig. 1.a). We used glass coverslips with a refractive index of n_2_= 1.51 and a phantom sample with a refractive index adjusted to n_3_= 1.38 using glycerol to approximate brain tissue in terms of refractive index [39]. As the beads are sitting on the bottom side of the coverslips, no aberration originates from the phantom sample. Nevertheless, we adjusted the refractive index to obtain a similar amount of reflected light at the glass-sample interface as is present during *in vivo* brain imaging. Using these samples, we experimentally measured the effective PSF for two objectives with different immersion media commonly used for *in vivo* imaging, and performed the corresponding numerical simulations. We used both a water (n_1_= 1.33) (LUMFLN 60x, NA1.1, Olympus) and a silicone oil (n_1_= 1.42) immersion objective (UPLanSAPO, 60x, NA1.3, Olympus). Figure 3 shows a quantitative analysis of the impact of the tilt angle on the fluorescence signal (maximal intensity in the PSF - left column) as well as on lateral and axial spatial resolution (middle and right column, respectively). In all graphs, markers with error bars represent experimental data, while the smooth lines show the results of the numerical simulations.

Figure 3 shows that the 2P signal is strongly affected by the tilt angle. Indeed, using a silicone oil objective (Fig. 3 – top line) only about 25% of the signal remains at a tilt angle of 1.5°. Tilt angles of this magnitude can easily be reached in case of *in vivo* imaging, but not necessarily spotted by eye, if no measures are taken to minimize the angle between 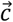 and 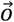, especially when looking at brain regions located on the side of the brain (e.g., cerebellum) or in depth (e.g., hippocampus). The effect on the resolution is less critical but still noticeable: the FWHMs of both lateral and axial fluorescent spots, scale with increasing tilt angles. Switching on the STED beam still improves the resolution, compared to the 2P case, even at relatively large tilt angles. Yet, at 1.5° tilt, a loss of at least 45% in lateral and 120% in axial resolution are observed compared to the case of orthogonal coverslips. The effect becomes even more evident when using a water immersion lens, where the gain in resolution almost completely vanishes at 1.5°. This results from the stronger refractive index mismatch at the immersion medium-glass interface, when using water instead of silicone oil as immersion medium.

**Fig. 3.**
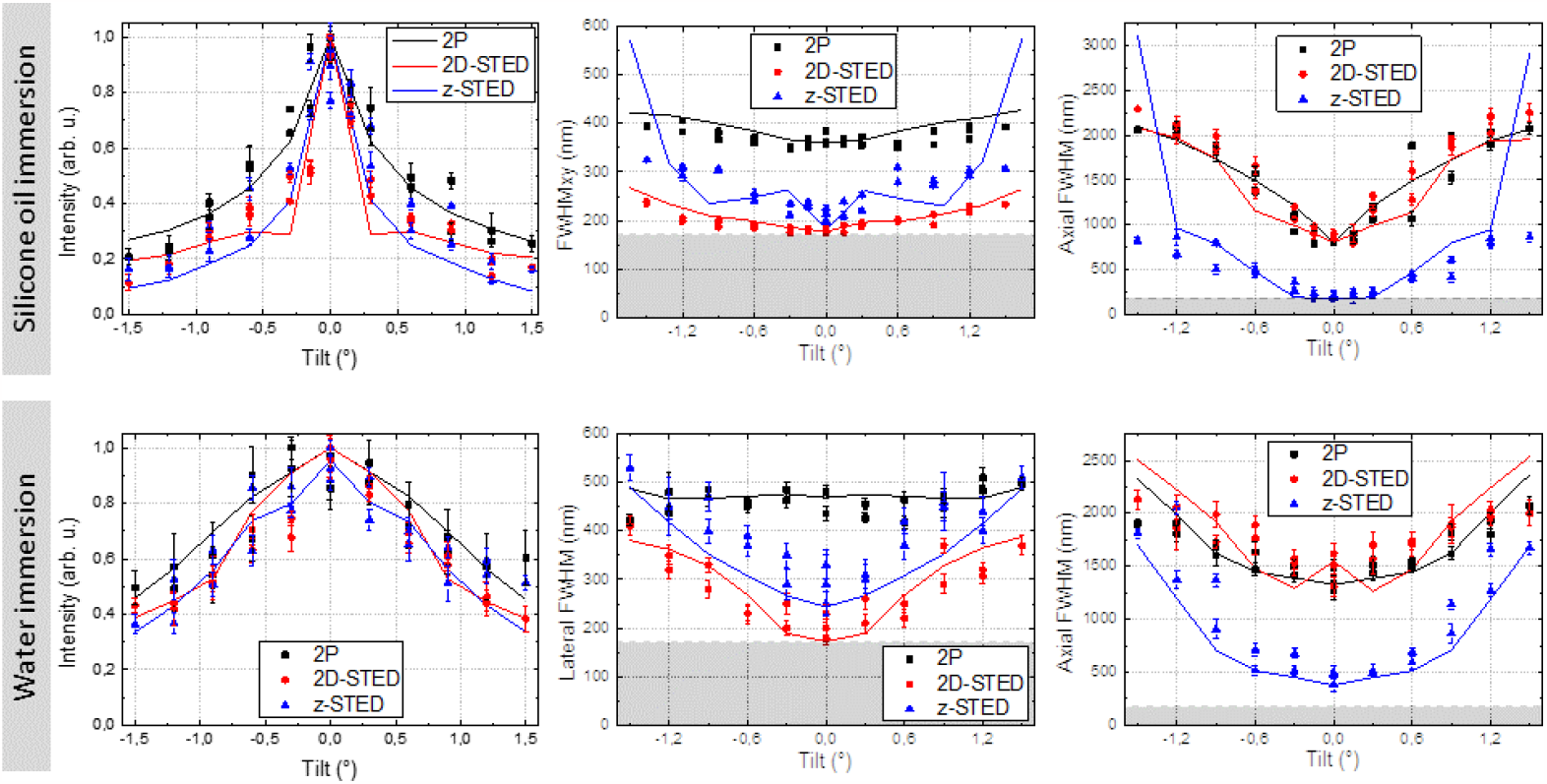
Quantification of the impact of the tilt angle of the coverslip on the fluorescence signal intensity (left column) and on the resulting lateral (middle column) and axial resolutions (right column) for silicone oil (top row) and water immersion lens (bottom row). Markers indicate individual measurements, with uncertainty as error bars. Straight lines represent the numerical calculation. Grey rectangles indicate the limit due to the physical size of the beads used in these experiments (170 μm in diameter).

### 3.2 Coverslip tilt measurement and correction

Having characterized the effect of a tilted coverslip on the microscope PSF, we then set out to propose a straightforward and robust approach to measure and correct it. So far only a few approaches have been proposed in the literature to deal with this issue in the context of imaging systems.

Theoretically, the use of a profilometer would enable accurate measurements of the tilt and flatness of the surface [40]. However, as the real concern is not the horizontality of the coverslip but its orthogonality to the optical axis of the objective lens, one would need to characterize the surface through the microscope objective. This is highly impractical, not to mention that profilometer are rarely available in life science labs. Recently, Galiñanes *et al*. [41] developed a very accurate approach to measure the tilt, however it requires a complex add-on device to be adapted and rotated directly on the objective. Alternatively, Steffens *et al*. [20] used an effective, but tedious, *pre hoc* alignment procedure for *in vivo* STED imaging in the mouse brain.

Our goal here was to introduce a procedure of alignment that could be used routinely with every animal used for *in vivo* STED imaging. It is worth noting that beyond STED imaging, considering the critical impact of the tilt on 2P signal (see Fig. 3), such an approach would be useful more generally for *in vivo* multiphoton imaging.

In our microscope, one of the simplest ways to measure the tilt is to perform an XZ scan (Fig. 4.a) of the coverslip-sample interface, while measuring the fluorescence from the sample. We evaluated the accuracy of this approach by introducing a controlled tilt angle, using the stage. Although this approach is quite efficient when imaging a fluorescent solution (calcein 100 μM) (Fig. 4.b), it performs less well on more realistic phantom samples, formed by a sparse ensemble of fluorescent beads attached to the coverslip (Fig. 4.c). Indeed, the uneven distribution of bright structures leads to a significant discrepancy between the introduced tilt versus the measured tilt angle (Fig. 4.d). In addition, to apply this approach *in vivo*, it would be necessary to place fiducials on the coverslips, prior of implanting the cranial window, which represents an additional challenge for *in vivo* imaging and might result in out of focus fluorescence.

**Fig. 4.**
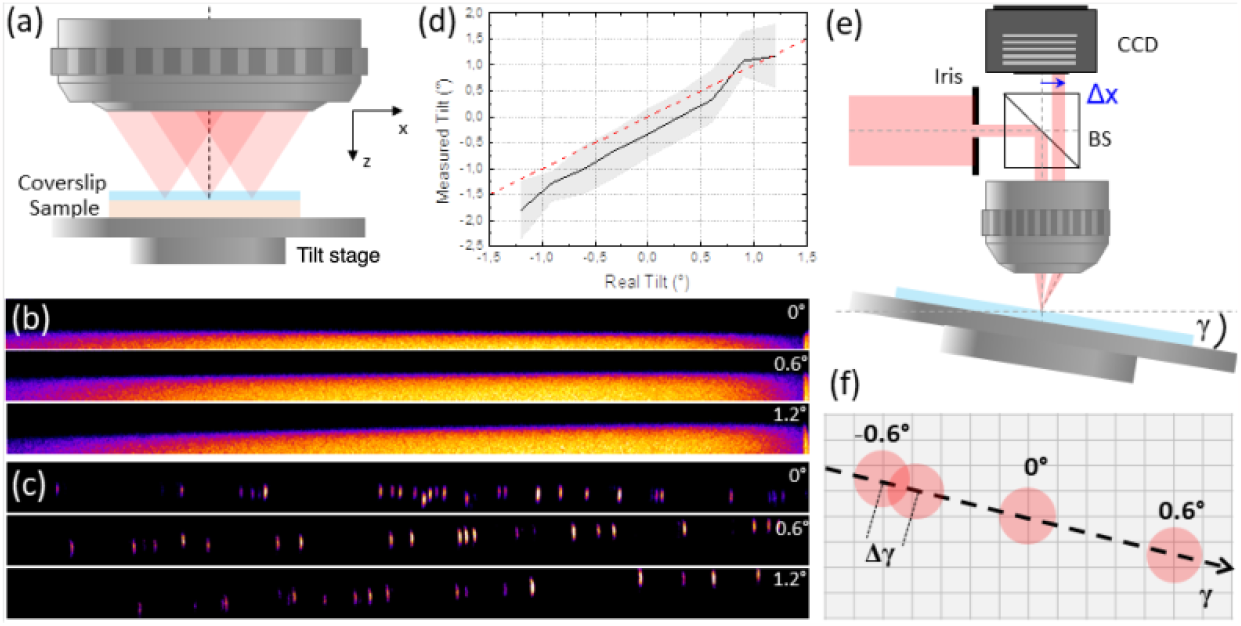
(a) Schematic of the XZ scan configuration used to determine the horizontality of the coverslip. (b) XZ images of a calcein solution covered with a coverslip, at different tilt angle. Image size: 150×10 μm^2^. (c) Fluorescent beads imaged through a coverslip at different tilt angle. Image size: 150×10 μm^2^. (d) Extraction of the tilt angle from the XZ scan as a function of the known tilt angle introduced. This highlights the relatively low accuracy of this approach. Black line indicates the measured tilt within confidence interval. Red dotted line indicates the expected curve. (e) Schematic of the setup added to the microscope to determine the tilt angle. (f) Schematic of the procedure used to determine the tilt from the displacement of the reflected spot recorded on the CCD camera.

To overcome this limitation, we developed an alternative approach based on a simple add-on device to our microscope and the reflection of the light on the coverslip. (Fig. 4.e). An iris and a 50/50 non-polarizing beam splitter were moved into the beam path, in place of the routing mirror between the tube lens and the objective. After passing through the objective, the laser beam is reflected on the coverslip surface, collected back through the objective with half of the signal passing via the beam splitter to the camera (Infinity 2). Closing the iris decreases the spot size on the back pupil of the objective and hence on the camera. Note that under-filling the back aperture of the objective leads to loose focusing, which makes the spot on the camera less sensitive to the axial position of the coverslip in the focus. Using the stage we introduced a controlled tilt angle and measured the displacement of the reflection spot on the camera (Fig. 4.f, Supp. Fig. 4 and Supp. Movie).

In this configuration, the precision of the tilt angle is limited by the ability to distinguish two positions of the reflected spot on the camera (see Fig. 4.f). This can be estimated by the width of the emission PSF on the CCD camera, which is itself given by the size of the iris (200 μm). The tilt angle Δγ precision is then given by:

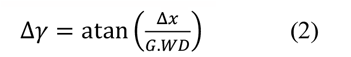

where Δx is the FWHM of the reflected spot on the CCD and WD (resp. G) is the working distance (the magnification, respectively) of the objective. Therefore, this procedure allows for a very precise determination of the coverslip tilt angle down to a precision of 0.07°. Importantly, this high precision largely derives from the use of a long working distance objective lens, as commonly used for *in vivo* imaging. Note that we focused our analysis here on the reflection from the immersion medium-glass interface, since this corresponds to the highest refractive index mismatch when using a water objective. Yet, for oil objectives, it would be possible to adapt this approach to exploit the reflection on the glass-sample interface.

Alternatively, in the context of STED microscopy, it is possible to determine the optical aberrations to retrieve the inclination of the coverslip. Using a spatial light modulator or deformable mirror, one could iteratively optimize the PSF by looking at the reflection from gold beads. Note that such approaches could be automated using a feedback loop from the microscope output and optical modeling or machine learning. This approach would provide a calibration of the aberration introduced by the coverslip (Supp. Fig. 5), which could then be used retrieve the tilt angle. However, this approach would only be efficient in the context of a shaped PSF, such as *donut* and *bottle* beams, in which the distortion introduced by the tilted coverslip are particularly obvious.

## 4. Conclusion

In this work, we numerically and experimentally investigated the effect of a tilted coverslip on 2P and STED microscopy, using water and silicone oil immersion objectives. While the effective resolution becomes substantially degraded with tilt angles on the order of 1°, the effect on the intensity of the signal is even more pronounced. We then propose and compare a solution to measure and correct the tilt angle directly below the microscope objective using a simple device added to the microscope. This will facilitate the optimization of super-resolution imaging, and more generally of multiphoton microscopy, especially applied to live animals.

## Acknowledgements

This project received funding from the European Union’s Horizon 2020 research and innovation program under the Marie Skłodowska-Curie action (Grant No. 794492), the Fonds AXA pour la Recherche to SB, the Doctoral School for Health and Life Sciences of the University of Bordeaux to JR, the European Research Council (ERC-SyG ENSEMBLE) (Grant No. 951294), Human Frontier Science Program (Grant No. RGP0036/2020), ERA-NET NEURON (Grant No. ANR-17-NEU3-0005), the Fédération pour la recherche sur le cerveau (FRC), and Agence Nationale de la Recherche (Grant No. ANR-17-CE37-0011) to UVN.

## Data availability statement

All data, code, and materials used in the analysis are available for non-commercial research purposes upon reasonable request.

## Supplemental documents

See Supplemental movie and figures for supporting content.

