## Supplementary figures and images for "Impact of a tilted coverslip on two-photon and STED microscopy"

### Supplemental figures

#### Supplemental figures

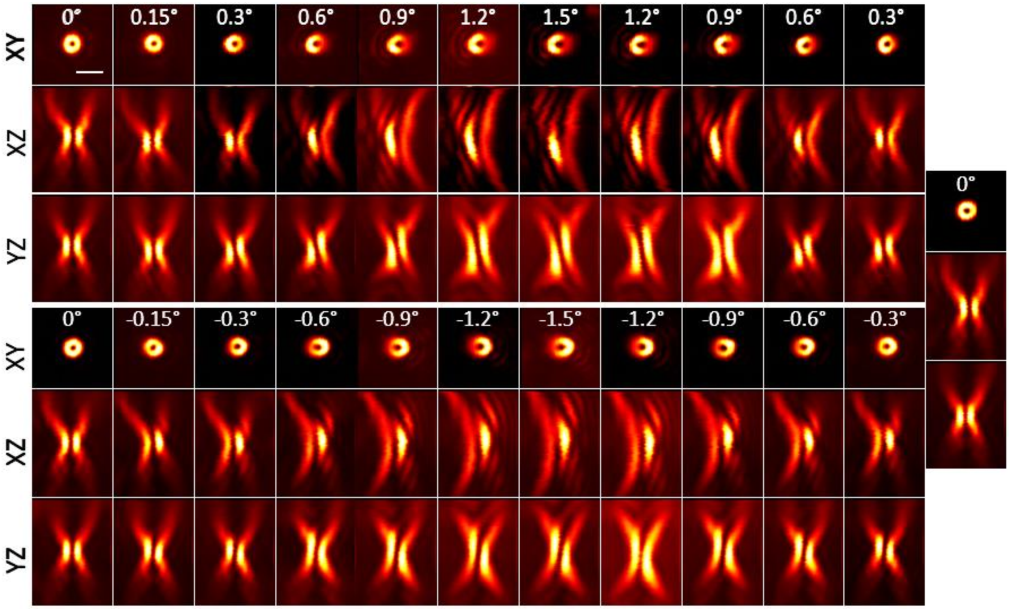


**Figure S1**

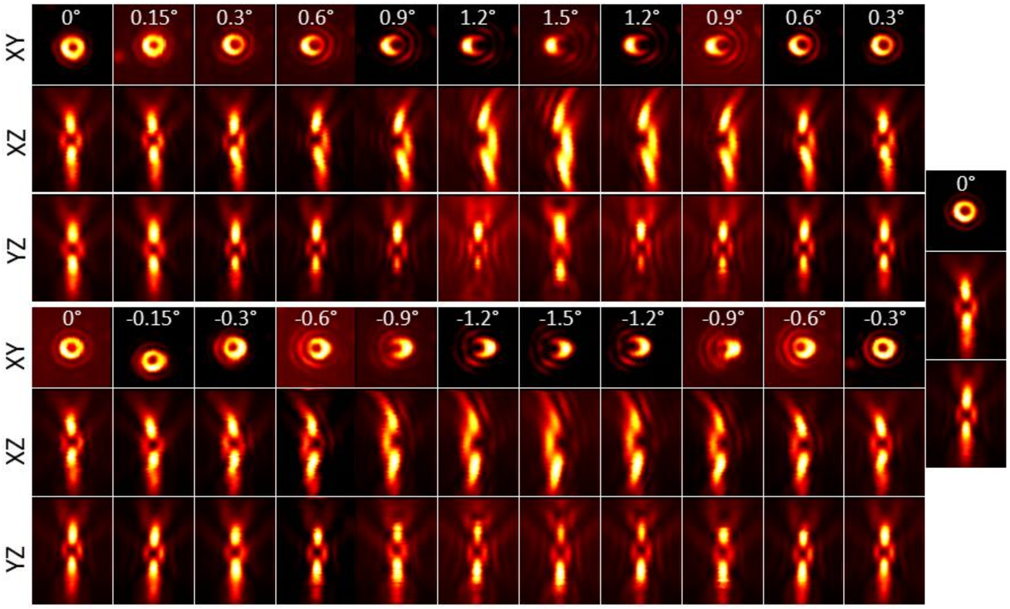


**Figure S2**

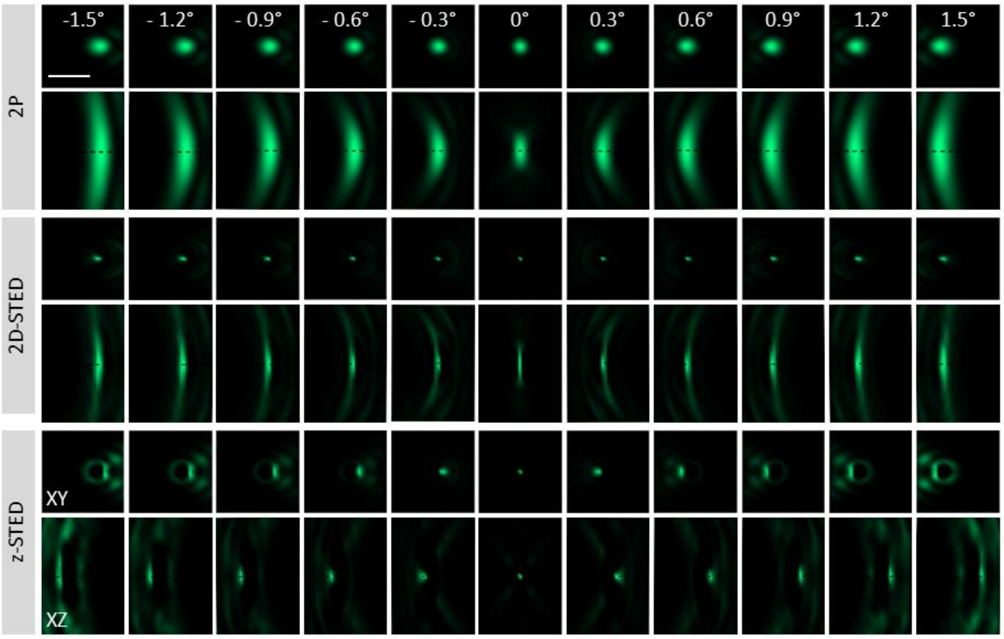


**Figure S3**

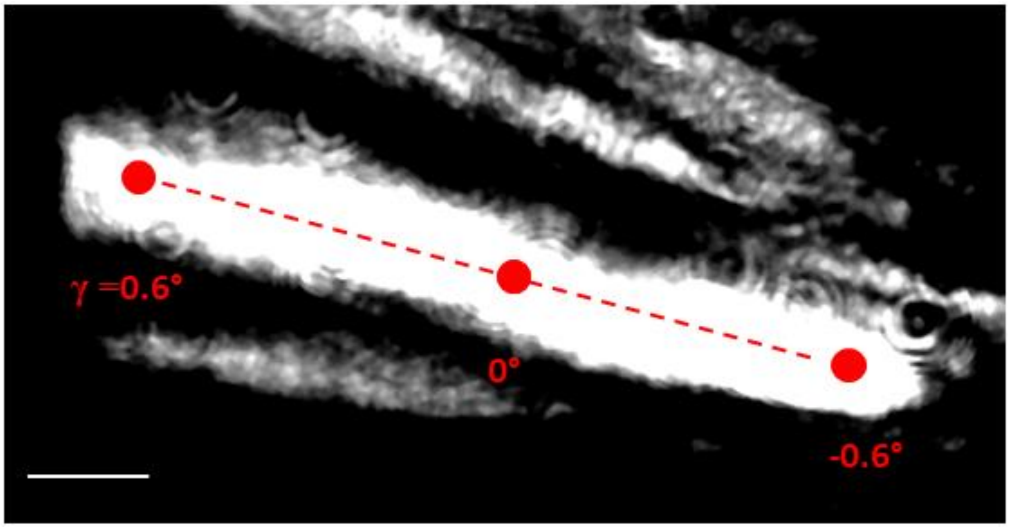


**Figure S4**

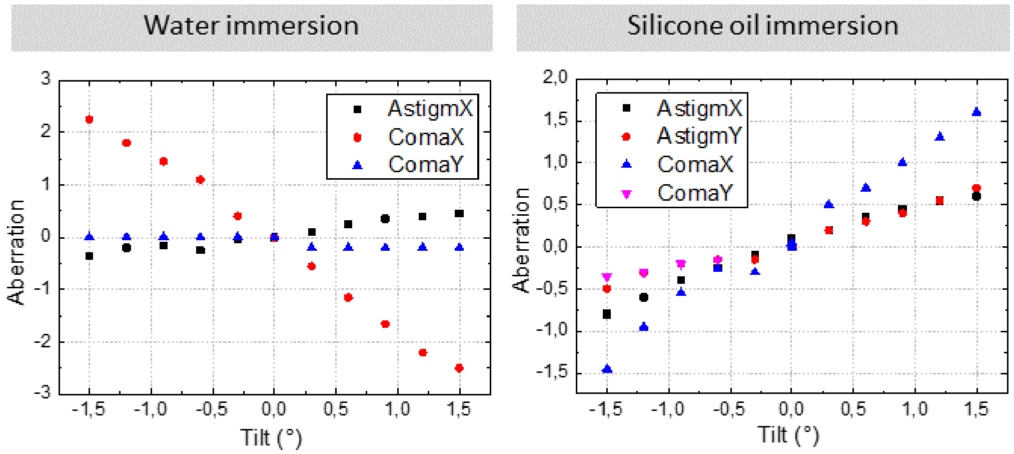
